# Quality versus quantity: The balance between egg and clutch size among Australian amphibians is related to life history and environmental conditions

**DOI:** 10.1101/2020.03.15.992495

**Authors:** John Gould, Chad Beranek, Jose Valdez, Michael Mahony

## Abstract

An inverse relationship between egg and clutch size has been found repeatedly across animal groups, including birds, reptiles and amphibians, and is considered to be a result of resource limitations and physical constraints on the reproducing female. However, few studies have contextualised this relationship with respect to various environmental selecting pressures and life history traits that have also likely influenced the selection of an optimal egg/clutch size combination, while even fewer have tested these interrelationships using robust natural history datasets. In this study, we aimed to test current hypothesises regarding these relationships on both egg and clutch sizes among the Australian Anurans, which to date have not received this kind of investigation. Specifically, we looked at the influence of environmental selecting pressures (egg laying location, environment persistence and bioregion) and life history traits (adult female body size, egg development type, parental care level, breeding period and temporal breeding pattern). As expected, a clear inverse relationship was found between egg and clutch size, while female body size was positively related to both. Generally speaking, smaller clutches of larger eggs tended to be produced by species that i) oviposit terrestrially, ii) showcase direct development and iii) possess high levels of parental care. Temporal breeding pattern was strongly related to clutch size only, with large clutches occurring in explosive breeding species, while breeding habitat was strongly related to egg size only, with large eggs sizes occurring in terrestrial species. Altogether, these findings indicate that numerous factors have likely influenced the evolution of an optimal clutch type in this group, highlighting the importance of incorporating such variables into animal studies on egg and clutch sizes.

## Introduction

Considerable research has been conducted on the evolution of life-history patterns, which has been contextualised by several core principles relating to resource limitations (Lack 1954; Williams 1966) and r-K selection theory (Pianka 1970). The latter of these predicts that an organism will possess life history trait combinations that exist along a continuum defined by two extremes; a small investment of resources per propagule, large clutch size, little parental care and a short life span (r-type selection) or the opposite (K-type selection). It is now understood that there are a variety of environmental factors that influence life history, (Haywood & Perrins 1992; Adolph & Porter 1993; Murphy 1968; Stearns 1976), many of which have formed trade-offs in reproductive strategies that are directly related to propagule quantity and size. However, for many species groups, the environmental context in which reproduction occurs and the subsequent relationships between these aspects of reproduction remain to be tested using robust natural history datasets.

The partitioning of a female’s finite resources across her reproductive life defines how a species has adapted to the selecting pressure of the environment in which it resides and will have considerable influence on parent and offspring fitness (Stearns 1976). As finite resources expended on current reproduction will reduce potential survival and, as such, additional chances to reproduce (Williams 1966), a trade-off must be made between the total resources invested per clutch and the number of clutches produced over reproductive life (i.e. semelparity versus iteroparity) (Hughes 2017). Further, a trade-off must also be made between the number of eggs produced per clutch and their size, given a female’s finite egg-carrying capacity and reproductive reserve (Lack 1967). These constraints will generally result in an inverse relationship between egg size and number per clutch (Parker & Begon 1986; Dziminski *et al.* 2009), with selection favouring either the production of large eggs (provided there are few) or many eggs (provided they are small). When considered from the perspective of r-K selection theory, this is a trade-off between offspring quality (amount of resources invested per offspring) and quantity (productivity via the number of propagules produced). As such, a conflict between maternal fitness and offspring fitness arises (Trivers 1974), given that large offspring are generally fitter (Pianka 1970; Moran & Emlet 2001; Einum & Fleming 1999) but will come at the expense of fecundity. Selection for a particular clutch type will be influenced by the effect of egg size on offspring survival under a given set of environmental conditions, along with the influence of various other traits that make up a specie’s life history, which is assumed to result in the evolution of an optimal egg size (Smith & Fretwell 1974; Lloyd 1987; Roff 1992; Einum & Fleming 2004). In variable environments, it may not be possible for an optimal egg size to evolve, with greater fitness benefits obtained by selecting for the deliberate production of variable egg and clutch sizes through forms of adaptive plasticity when reliable cues to the future state of the environment are available (Fox *et al.* 1997) or bet-hedging when they are not (Olofsson *et al.* 2009; Einum & Fleming 2004). However, examining reproductive investment within the framework of optimal egg size theory may provide a wealth of information on the general influence of these factors within species groups.

An inverse relationship between egg size and number per clutch has been recorded in several animal groups (Williams 2001; Herman & Bout 1998). Among the amphibian, this relationship has been shown repeatedly and verified in several studies (Duellman 1989; Crump 1974; Crump & Kaplan 1979; Perotti 1997; Hödl 1990). While a majority of amphibians are aquatic breeders and exhibit mostly r-type traits, including the production of small eggs and provision of limited care, there is a continuum of egg and clutch sizes, as well as a diverse array of reproductive patterns that occur in this group (Duellman & Trueb 1986; Salthe & Duellman 1973; Haddad & Prado 2005). Some species have become entirely independent of the need of an aquatic environment for embryonic and larval development altogether, laying eggs on land that undergo direct development, while others have an intermediate mode. For example, the red-crowned toadlet (*Pseudophryne australis*) has an intermediate mode of reproduction where the eggs are laid in a terrestrial nest and develop to an advanced stage before hatching into tadpoles (Thumm & Mahony 2002). Amphibians also exhibit a wide range of forms of parental care, from egg attendance through to energetically expensive activities, such as offspring feeding and movement (Furness & Capellini 2019; Crump 1996).(Zamudio *et al.* 2016). Despite these differences, an overwhelming majority of amphibians are oviparous. As such, factors that have likely influenced the evolution of a particular egg and clutch size trade-off will primarily include the environmental conditions offspring are exposed to throughout embryogenesis and later stages of development.

Several theories and evolutionary patterns have been proposed to explain the influence of key factors on the relationship between egg and clutch size in anurans based on their influence on offspring survival. Differences are expected to occur between species that exploit different egg laying environments (terrestrial versus aquatic) and their variability (temporary versus permanent), as this will dictate exposure to risk factors such as competition, predation, sub-optimal temperature and moisture levels (Hendry *et al.* 2001; Werner & McPeek 1994; Skelly *et al.* 1999; Wilbur 1987; Babbitt *et al.* 2003; Semlitsch *et al.* 2015; Skelly 1996; Wellborn *et al.* 1996). For example, the high risk of desiccation in temporary aquatic systems due to short hydroperiods should select for the production of smaller eggs, as this will result in shorter developmental periods and an improved chance of offspring metamorphing prior to pond dry-up (Salthe & Duellman 1973; Kaplan 1980; Brown 1989). In contrast, the production of larger eggs in terrestrial egg laying species may improve their resistance to evaporative water loss and/or facilitate direct development so that the absence of free-standing water is not a limiting factor for offspring survival at all (Bogart, 1981, Roberts, 1981, Bradford and Seymour, 1988). Underpinning exposure to environmental risk factors will be the physical location in which reproduction occurs. For example, Duellman (1989) argues that terrestrial reproductive modes can only evolve in humid environments such as tropical forests due to the otherwise increased risk of egg dehydration, which should lead to a discrepancy in egg sizes between bioregions. The level of parental care that is provided to offspring will also be an influencing factor as it will determine the level of exposure to an external environment and possible suboptimal conditions (Furness & Capellini 2019; Summers *et al.* 2005). Other behavioural traits pertaining to life history, such as the length of the breeding period and whether breeding is explosive or prolonged, will also influence exposure to risk factors or possibly even result in their own risks to offspring survival (Petranka & Thomas 1995; Morin *et al.* 1990). In combination, it is these aspects to reproduction that will be primary contributors in the evolutionary shaping of an optimal clutch.

Among the Australian anurans there has been little examination of the relationship between egg and clutch size. This is despite them being an ideal model for this investigation given the large number of species in this group. Further, a large proportion of these species are derived mainly from two large and likely monophyletic lineages (Hylidae and Myobatrachidae) with a long evolutionary history of isolation on the continent (Heyer & Liem 1976; Farris *et al.* 1982; Hutchinson & Maxson 1987; Tyler 1971), with more recent invasions by Microhylidae and Ranidae from Asia (Tyler 1989). This is favourable for analysing the influence of the environment on reproduction, as the reproductive patterns currently found among species in these families likely represents the radiation of a few common ancestral genotypes into various ecological niches.

In light of this, we aimed to test current hypothesises regarding known reproductive trade-offs, as well as environmental selecting pressures, on the evolution of egg and clutch size in this group. We collated natural history data pertaining to the reproduction of 128 Australian anurans from three endemic families with a long evolutionary history on the continent (Hylidae, Myobatrachidae, *and* Limnodynastidae), as well as for one family which is a relatively newer endemic (Microhylidae). This constitutes 53% of the currently described Australian amphibian species. This data was used to investigate whether i) mean egg size is inversely related to mean clutch size, ii) egg and clutch size are related to female body size, and iii) egg and clutch size are related to environmental selecting pressures (egg laying location, environment persistence and region) and life history traits (development type, parental care level, breeding period and temporal breeding pattern).

## Methods

### Data collection

The partitioning of clutch resource between egg size and number, along with factors pertaining to reproductive pattern, were identified for each species using data collated by Anstis (2017). This seminal work by Anstis (2017) is the most complete and up to date dataset on the reproductive biology of Australian anurans, including original work and more than 230 references, providing an opportunity by which to compare reproductive patterns between species in this group. Missing data values were obtained from online databases (AmphibiaWeb 2018; Oliveira *et al.* 2017) when they were not available in Anstis (2017). To enable future comparisons to be conducted using the same methodology, we have provided precise definitions for each variable. Table 1 contains an overview of the variables collected for each species.

**Table 1.**
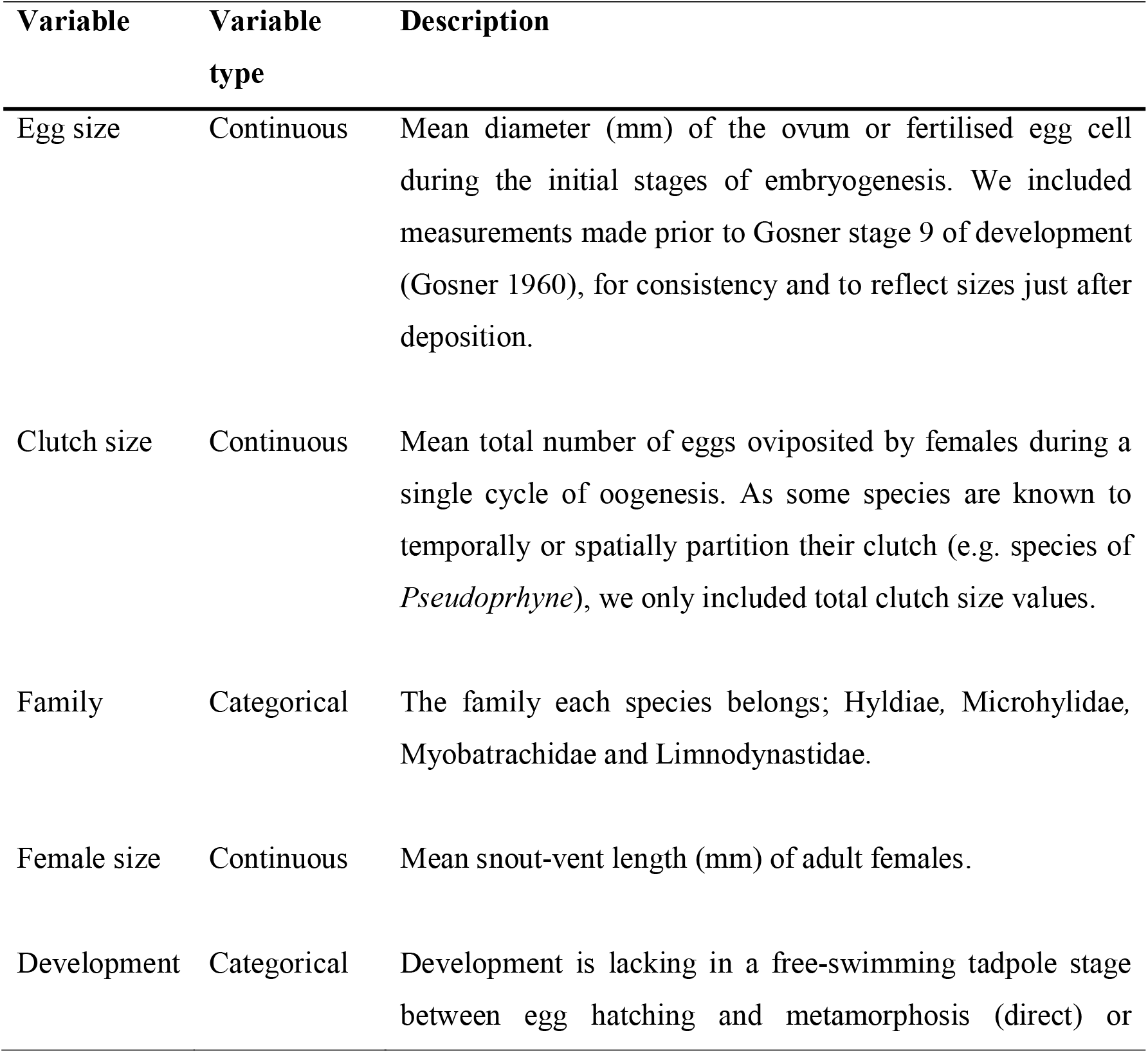
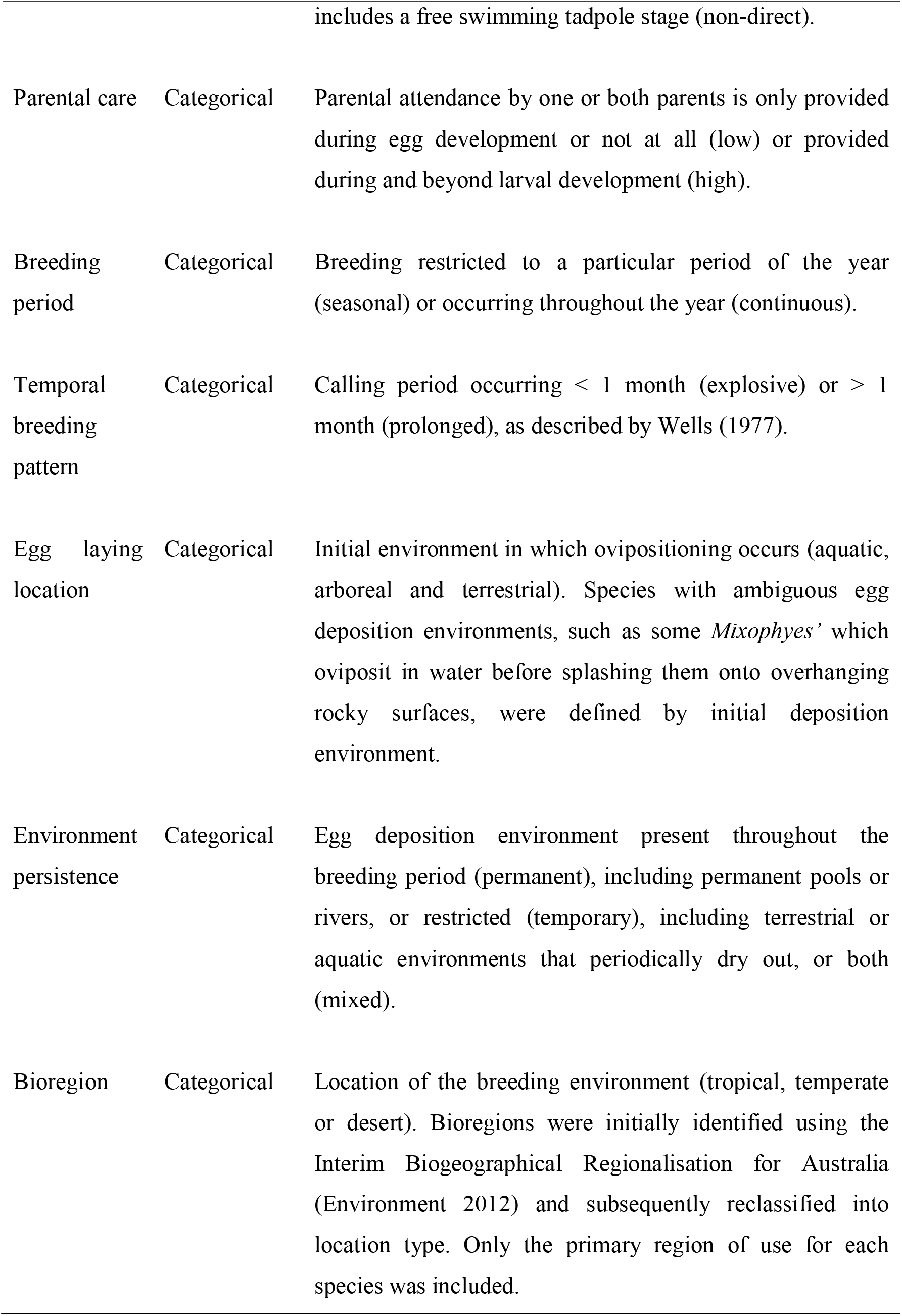
Reproductive variables obtained from Anstis (2017) for Australian anurans. Definitions for variables have been provided.

| Variable | Variable type | Description |
| --- | --- | --- |
| Egg size | Continuous | Mean diameter (mm) of the ovum or fertilised egg cell during the initial stages of embryogenesis. We included measurements made prior to Gosner stage 9 of development (Gosner 1960), for consistency and to reflect sizes just after deposition. |
| Clutch size | Continuous | Mean total number of eggs oviposited by females during a single cycle of oogenesis. As some species are known to temporally or spatially partition their clutch (e.g. species of *Pseudophryne*), we only included total clutch size values. |
| Family | Categorical | The family each species belongs; Hyldiae, Microhylidae, Myobatrachidae and Limnodynastidae. |
| Female size | Continuous | Mean snout-vent length (mm) of adult females. |
| Development | Categorical | Development is lacking in a free-swimming tadpole stage between egg hatching and metamorphosis (direct) or |
|  |  | includes a free swimming tadpole stage (non-direct). |
| Parental care | Categorical | Parental attendance by one or both parents is only provided during egg development or not at all (low) or provided during and beyond larval development (high). |
| Breeding period | Categorical | Breeding restricted to a particular period of the year (seasonal) or occurring throughout the year (continuous). |
| Temporal breeding pattern | Categorical | Calling period occurring < 1 month (explosive) or > 1 month (prolonged), as described by Wells (1977). |
| Egg laying location | Categorical | Initial environment in which ovipositioning occurs (aquatic, arboreal and terrestrial). Species with ambiguous egg deposition environments, such as some *Mixophyes* which oviposit in water before splashing them onto overhanging rocky surfaces, were defined by initial deposition environment. |
| Environment persistence | Categorical | Egg deposition environment present throughout the breeding period (permanent), including permanent pools or rivers, or restricted (temporary), including terrestrial or aquatic environments that periodically dry out, or both (mixed). |
| Bioregion | Categorical | Location of the breeding environment (tropical, temperate or desert). Bioregions were initially identified using the Interim Biogeographical Regionalisation for Australia (Environment 2012) and subsequently reclassified into location type. Only the primary region of use for each species was included. |

It must be recognised that sample sizes over which measurements of egg, clutch and female size were calculated differed between species, reflecting limitations in natural history data. Even though various factors are known to cause interspecific variations in egg and clutch size, such as female size and condition (Long 1987; Gibbons & McCarthy 1986), we assumed that mean estimates were representative for each species. We also assumed that estimates reflect an optimal partitioning of within-clutch resources for each species, though it is possible that some species may show within-female variations that is adaptive bet-hedging or plasticity (Bernardo 1996; Lips 2001).

### Statistical analyses

We investigated whether there was an inverse relationship between egg and clutch size among the Australian anurans using simple linear regression, along with the Pearson Correlation Coefficient. Mean egg size was compared to mean clutch size, both before and after adjustment by adult female size. The partitioning of clutch resources was subsequently analysed using generalised linear mixed modelling. Egg size and clutch size were analysed as dependent variables in separate models, with the predictors *female size, development type, parental care, breeding period, temporal breeding pattern, egg laying location, environmental persistence, and bioregion* included as fixed effects, and *family* included as a random effect. All continuous variables, including *egg size*, *clutch size* and *female size* were log transformed prior to analysis. Effect sizes of the full models, along with 95 % confidence intervals, were used as they best reflect the range of predictors investigated, decrease bias in the predicted values, and ensure a balanced representation of non-significant results compared to high false detection rates, error rates, and biases associated with model selection (Hegyi & Laczi 2015; Mundry & Nunn 2009; Forstmeier & Schielzeth 2011; Whittingham *et al.* 2006; Flom & Cassell 2007). All modelling was performed using the RStudio statistical software package (Team 2015).

## Results

A complete dataset was obtained for 128 species, representing approximately 56% of all Australian anurans. Between species, there were considerable differences in mean egg size (0.8 to 5.1 mm), mean clutch size (6 to 8000 eggs), and mean female size (16.6 to 135 mm). Species were mostly from the families Myobatrachidae (36%), Hylidae (33%) and Limnodynastidae (24%), with only a small number from Microhylidae (7%). A majority of species were prolonged breeders (83%), possessed non-direct development (98%), and exploited environments that were aquatic (71%) and temporary (53%). Most species showed seasonal breeding (85%) and low levels of parental care (98%). Most species were found in temperate or tropical regions (54% and 41%, respectively), with few species found in deserts (5%).

### The relationship between egg and clutch size

Egg size was inversely related to log clutch size (r = −0.6072, t value = −8.578, p < 0.0001) with an R^2^ of 0.37. Each 1 mm increase in egg size resulted in the reduction of clutch size by 0.25 % (Fig. 1). A stronger relationship was found when egg and clutch size were adjusted for by female size (r = −0.8501, t value = −18.12, p < 0.0001), with an R^2^ of 0.7227 (Fig. 1).

**Figure 1.**
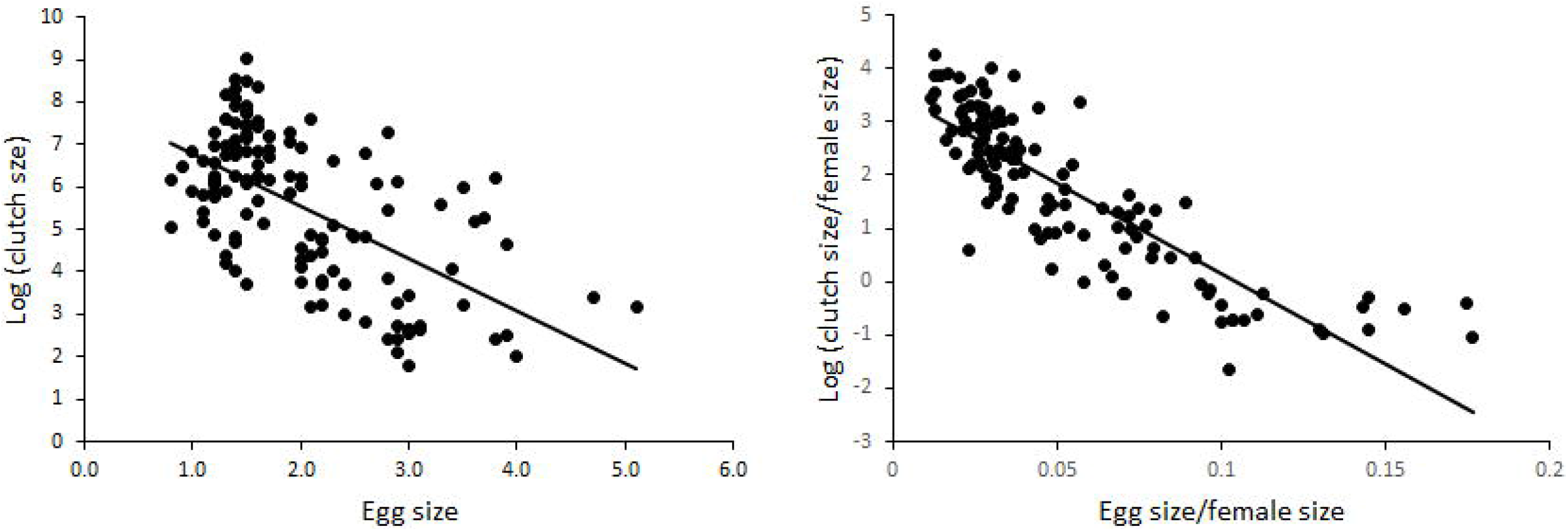
Inverse relationship between egg and clutch size among Australian anurans. Mean egg and clutch values were obtained for 128 species. This relationship has been presented before (left) and after (right) adjustment for by female size. Clutch size has been log transformed.

### Influence of egg size

The saturated multiple regression comparing egg size to all eight predictors had a conditional R^2^ of 0.70. The log of mean *egg size* was positively correlated with the log of mean *female size* (F value = 29.0725, p < 0.0001), with a 1 % increase in female size resulting in a 0.36 % increase in egg size. The smallest eggs (diameter = 0.8 mm) were produced by *Litoria Microhylidaebelos*, while the largest eggs (diameter = 5.1 mm) were produced by *Myobatrachus gouldii*. Categorical variables that were found to be related to egg size included *development type* (F value = 4.3290, p = 0.0417), *parental care level* (F value = 12.5656, p = 0.0006), *environmental persistence* (F value = 5.1508, p = 0.0072), and *egg laying location* (F value = 25.3798, p < 0.0001). Based on pairwise comparisons between levels of each categorical variable, greater egg sizes were found for species which were direct developers (z value = 2.081, p = 0.0375), with high levels of parental care (z value = 3.545, p = 0.0004), those which exploited permanent compared to temporary breeding environments (z value = 3.105, p = 0.0054), and for terrestrial (z value = 6.567, p < 0.001) and arboreal breeders (z value = 3.591, p < 0.001) (Fig. 2a).

**Figure 2.**
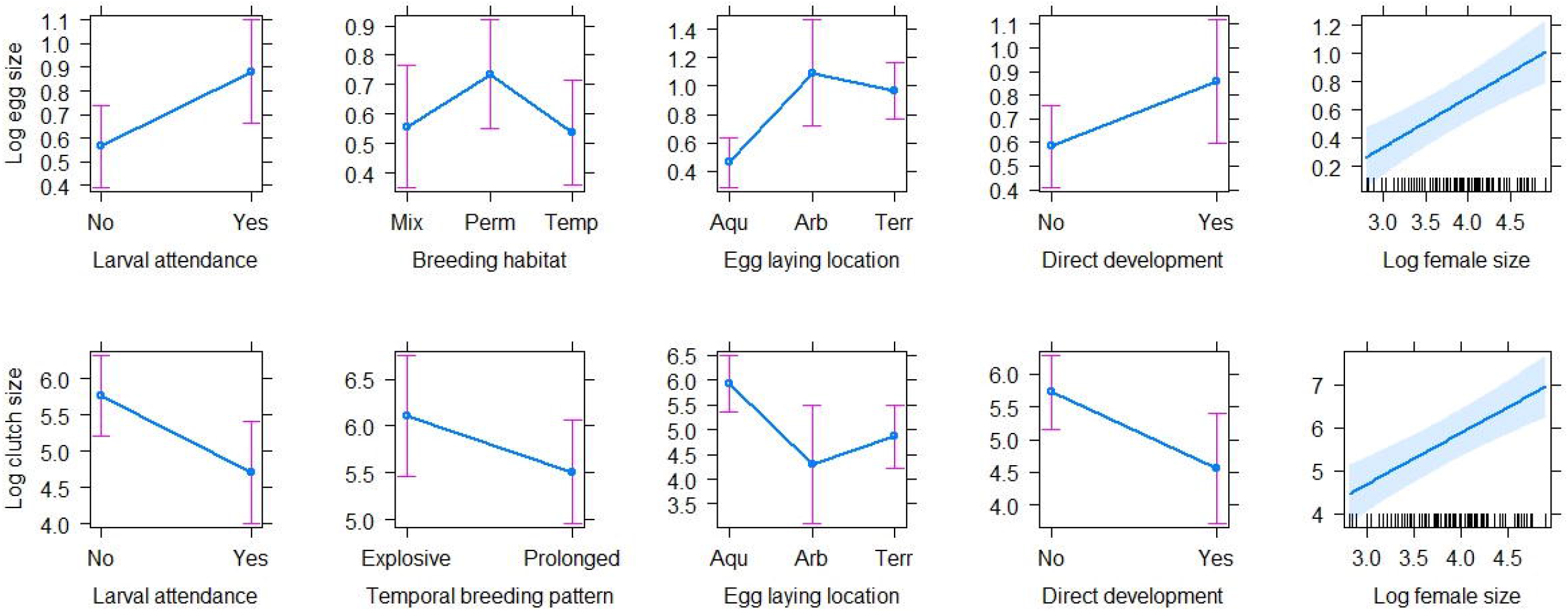
The relationship between environmental variables and life history traits on the partitioning of clutch energy among Australian anurans. Shown are significant predictors following mixed modelling with log mean egg size (a) and log mean clutch size (b) as the dependent variable. Differences in mean effect size of levels for each significant categorical variable have been displayed with 95 % confidence intervals. Data was obtained for 128 species.

### Influences of clutch size

The saturated multiple regression comparing clutch size to all eight predictors had a conditional R^2^ of 0.81. The log of mean *clutch size* was positively correlated with the log of mean *female size* (F value = 31.5787, p < 0.0001), with a 1 % increase in female size resulting in a 1.19 % increase in clutch size. The largest female species examined, *Litoria splendida* (SVL = 118) and *Litori infrafenata* (SVL = 135 mm), possessed mean clutch sizes of over 8000 and 4000 eggs, respectively, compared to the smallest species, *Cophixalus hosmeri* (SVL = 17 mm) and *Geocrinia vitelline* (SVL = 18 mm), with mean clutch sizes of only 6 and 11 eggs, respectively. *Litoria aurea* had the second largest clutch size, and had the fifth largest female length (SVL = 108 mm). Categorical variables that were found to be related to clutch size included *development type* (F value = 8.5560, p = 0.0048), *breeding type* (F value = 6.3598, p = 0.01306), *parental care level* (F value = 13.9524, p= 0.0003), and *egg laying location* (F value = 12.7155, p < 0.0001). Based on pairwise comparisons between levels of each categorical variable, greater clutch sizes were found for non-direct developers (z value = 2.9257, p = 0.0034), explosive breeders (z value = 2.522, p = 0.0117), those with low levels of parental care (z value = 3.735, p = 0.0002), and for aquatic breeders compared to terrestrial breeders (z value = 4.438, p < 0.0010) and arboreal breeders (z value = 2.948, p = 0.0080) (Fig. 2b).

## Discussion

We demonstrate clear relationships between reproductive traits and life history and environmental variables for the first time among the Australian anurans using a large dataset. A clear inverse relationship was found between egg and clutch size among species in this group, mirroring results obtained in other amphibian groups (Duellman 1989; Crump 1974; Crump & Kaplan 1979; Perotti 1997; Hödl 1990), and supporting the known trade-off that must occur between the provision of resources per egg and the number of eggs produced per clutch (Parker & Begon 1986). Further, both egg and clutch size were found to be positively related to female size, highlighting the intimate link that exists between egg, clutch and female size in this group. A variety of life history traits and environmental variables were also related to the partitioning of clutch resources. Generally speaking, these variables were related to egg and clutch size in opposing manners, as expected given their strong inverse relationship. Smaller clutches of larger eggs tended to be produced by species that i) oviposit terrestrially, ii) have direct development and iii) possess high levels of parental care. Breeding type was strongly related to clutch size only, while breeding habitat was strongly related to egg size only. Altogether, these findings indicate that numerous environmental and life history factors are related to both egg and clutch size, suggesting that these factors have directly shaped the evolution of an optimal clutch among the Australian anurans.

### The relationship between egg, clutch and female size

Clutch size correlated positively with female size, with more eggs produced per clutch by comparatively larger species. This link between fecundity and size has been shown repeatedly in various amphibian assemblages (Berven 1988; Lemckert & Shine 1993; Prado & Haddad 2005), though this has not always been the case and some studies have not observed this pattern (Elmberg 1991; Tsiora & Kyriakopoulou-Sklavounou 2002). Even so, it comes as no surprise given that the maximum clutch size (in terms of volume) that an individual can produce will be constrained by the internal space (i.e. body cavity) available for ovarian development (Crump 1974). Although not tested as part of this study, it is possible that the evolution of a larger female size at sexual maturity may be an adaptive means of increasing the number of offspring produced per clutch without requiring eggs to be of a reduced size (Wells 2007, but see Shine 1988). No doubt, this accounts for sexual dimorphism in adult body size often seen in amphibians (Shine 1979; Shine 1989). On the other hand, growing to a large size may be particularly advantageous for species that have already reached a physiological limit in egg size (Sinervo & Licht 1991), or in circumstances where selection for a smaller egg size is not advantageous for offspring survival, though both hypothesise require testing.

Egg size was also positively related with female size, mirroring results obtained by Salthe (1969) in urodels. For some species, such as *Rana temporaria*, larger individuals have been shown to produce larger eggs (Gibbons & McCarthy 1986), which may be an allometric relationship that is the result of the scaling of organs associated with egg production (Schmidt-Nielsen & Knut 1984). However, there is little evidence in the literature to suggest that intraspecific differences in egg size are also morphologically constrained by a genetically predetermined adult size, as egg size should be selected to optimise offspring survival irrespective of female size. This is apparent among the Australian anurans, with some of the smallest species possessing some of the largest eggs. For example, *Austrochaperi pluvialis* has an SVL of 29 mm but the same egg size as that of *Mixophyes coggeri*, which has an SVL of 104 mm. Although we are cautious of these findings, this correlation does suggest that some level of variation in egg size among species is related to female body size, although this could be occurring in a manner that is not adaptive but simply an artefact of the scaling of body size.

### Temporal breeding pattern and environmental persistence

Larger clutches were recorded in species that exhibit explosive compared to prolonged breeding patterns, while smaller eggs were recorded by species that exploit temporary compared to permanent environments. An explosive breeding strategy is characteristic of many species that exploit temporary environments (Wells 1977; Crump 1974), such as ephemeral pools, where there are limited optimal periods for reproduction and/or restrictions on the availability of breeding sites throughout the season due to their hydrological regimes. In these environments, adults often must synchronise the onset of their reproduction with oncoming rainfall to maximise the chances of offspring development occurring when conditions are suitable (Reyer & Barandun 1997; Anholt *et al.* 1997). However, if these environments are relatively unpredictable, adults may be afforded only limited or unreliable cues to determine when suitable conditions arise (Moran 1992; Rungeand & Sherman 2002), leading to an increased risk that offspring will be exposed to suboptimal conditions if adults choose an incorrect time for egg deposition or if the length of these optimal periods is not sustained throughout offspring development. These factors may increase the uncertainty of offspring survival, leading to the selection for larger clutch sizes (of smaller eggs) to increase the odds of producing more breeding individuals in the subsequent generation (Salthe 1969).

The tendency for larger clutches of small eggs to be produced by species exploiting temporary environments could also be attributed to the relationship between egg size and development duration. When breeding sites are only available for short periods of time, it will be highly advantageous for adults to produce eggs which can complete development as quickly as possible, which may occur by reducing egg size (Salthe & Duellman 1973; Kaplan 1980). It is perhaps most noticeable for species which have come to exploit temporary waterbodies prone to repeated disturbance, where hydrological regime is the main driver of habitat quality and variability experienced by offspring (Lake 2000). Though smaller eggs will result in smaller offspring that may have less of a competitive ability and be more vulnerable to predation without alternative protective strategies (e.g. being poisonous) (Hayes *et al.* 2009; Travis 1980; Shine 1978), maternal fitness will be maximised given that offspring will be afforded a greater chance of being able to complete development and remove themselves from such short lived systems before they become incompatible with their continued development. As such, this represents a departure from the positive relationship expected between offspring size and performance, though such departures have been found in other animals (Moran & Emlet 2001; Kaplan 1992). This is apparent in *Lechridous fletcheri*, which produces in highly ephemeral pools that often dry up within a matter of days once rainfall has ceased and has a rapid larval developmental period of just 21 days (J. Gould, unpublished data, 2019). If, however, offspring survival is random with respect to size, the fitness benefits of producing larger eggs will still be nullified, and maternal fitness will be optimised by producing smaller eggs given the concomitant increase in fecundity (Morrongiello *et al.* 2012). As such, the benefits of producing small eggs in variable environments prone to disturbance, as well as those of decreasing environmental quality, may be two-fold; the small size of eggs may improve the chances of offspring developing more quickly while also affording females an ability to produce large clutches in systems where offspring survival is uncertain. Given this finding, we suggest that additional work needs to be carried out on the correlation between egg and metamorph size, and whether this size metric influences offspring developmental rate.

In contrast, the tendency for smaller clutches of large eggs to be produced in permanent systems is likely to occur as a result of the associated fitness benefits of a large offspring size (Pianka 1970; Moran & Emlet 2001; Einum & Fleming 1999), given that abiotic stressors in more stable systems of better quality will be weak relative to biotic interactions (Hendry *et al.* 2001). Likewise, the production of smaller clutches of large eggs may be favoured when breeding periods are prolonged, given that such breeding periods are more likely to be possible in permanent systems. However, prolonged breeding periods will also favour the production of smaller clutches in general so as to spread the risk of clutch failure over multiple breeding attempts within season (Griebeler *et al.* 2010).

### Parental care level

The occurrence of larvae attendance was correlated with the production of smaller clutches of larger eggs, which conforms to predictions from r-K type selection theory (Pianka 1970). Previous studies investigating the relationship between egg and clutch size in relation to parental care level in amphibians have resulted in varying findings, some which oppose r-K type selection theory (Nussbaum 1987) but a majority of which are in accordance (Vági *et al.* 2019; Summers *et al.* 2005; Summers *et al.* 2007). For example, in a study of the evolution of territoriality in frogs Vági *et al.* (2019) found that both large ovum size and small clutch sizes were strongly associated with extended parental care, which was found in an analysis of 700 species. Our results of sampled Australian anurans (56% of all Australian species) demonstrates that it may be more advantageous to produce fewer individuals that are larger with increasing levels of parental care. As such increased investment will reduce offspring exposure to environmental risk factors (Shine 1978), it will be more advantageous to invest more clutch resources per egg as the chances of survival will be comparatively much greater than for eggs that are left unattended. In order to facilitate increasing levels of care, however, there may be constraints on the number of offspring that such care can be adequately provided to (Cody 1966), which may also select for a smaller clutch. This may be particularly the case for the few Australian species that carry or brood their young, such as *Assa darlingtoni*, where there is limited space for such activities to be performed.

### Egg laying environment and direct development

As with terrestrial egg depositors, those which have evolved direct development tended to produce larger eggs in smaller clutches. The shared relationship between these two strategies and egg size among amphibians is of no coincidence, considering that direct development often occurs alongside terrestrial egg deposition (Günther 2006), with both removing many of the threats associated with having egg/tadpole stages reliant on free standing water to complete metamorphosis (Denver *et al.* 1998). Similar to eggs laid in ephemeral waters, those laid terrestrially will be vulnerable to changes in moisture levels, which may influence gas exchange or expose offspring to a risk of mortality as a result of desiccation (Bradford & Seymour 1988; Anstis *et al.* 2007). Producing larger eggs will buffer offspring from these threats, given that a smaller surface area relative to volume will reduce rates of evaporative water loss, thereby improving the longevity of the egg and the chances of the embryo successfully completing embryogenesis before internal water levels fall too low (Mitchell 2002; Salthe 1965; Bogart 1981; Roberts 1981; Bradford & Seymour 1988). However, it is apparent across amphibian groups that terrestrial development is confined to areas where moisture is predictable and stable, particularly in tropical rainforests, and where hygric stress is less likely to be a strong selecting pressure for the evolution of an optimal egg size.

Among the Australian anurans, relatively few species have evolved direct development (*M. gouldii*, both *Arenophryne*, all microhylids, and *Metacrinia nichollsi*), all of which are terrestrial egg depositors (Anstis 2017). In order for an egg to facilitate direct development, it must possess a comparatively larger supply of energy for the embryo to bypass the tadpole stage and spawn into a metamorph at the end of the developmental period (Summers *et al.* 2007). Presumably, this can only occur with an increase in the yolk content of the egg. Although direct development is found in species with different egg sizes, suggesting that an absolutely large egg is not necessary for direct development to be achieved, a relative increase in egg size has been found among urodels when species are grouped based on adult body size (Salthe 1969), suggesting the selection for a relative increase in egg size in direct developing species. Given these findings, the transition to terrestrial egg laying and direct development from the ancestral mode have likely both driven the selection for an enlarged egg size in tandem, albeit at the expense of a large clutch.

### Seasonal and region

Although physical micro-habitat location will underpin exposure to environmental risk factors that should directly influence the evolution of an appropriate egg size, it did not have an effect among Australian anurans in this study when analysed in terms of bioregion. Instead, it may indirectly have an effect by selecting for other traits which will alter the manner in which eggs are exposed to external risk factors. For example, while terrestrial breeding is considered to be confined to species inhabiting tropical environments, the Australian terrestrial breeder *M. gouldii* deposits its eggs in dry, semi-arid environments (Poynton 1964; Roberts 1984). To improve the suitability of the environment in which the eggs are exposed to, this species deposits in underground burrows that reach aquifers that will maintain moisture levels throughout development. As a result of this strategy, the incubation environment is not reflective of the true conditions offspring would be exposed to in that bioregion. Other terrestrial and direct developing species in this group, including *Arenophryne rotunda* and *M. nichollsi* also exploit temperate or arid environments, highlighting the possibility for such reproductive patterns to be selected for in areas far removed from the tropics on the caveat of there being some sort of reliable moisture source. Other geographical variables not considered as part of this study, such as altitude (Morrison & Hero 2003), are known to affect egg and clutch size among amphibian as well, though the highest mountains in Australia are less than 3000 m and there are only a few species considered to be alpine in distribution (e.g. *Pseudophryne corroboree* and *Philoria frosti*).

Likewise, breeding period had no effect on the partitioning of clutch energy. Given that the difference between seasonal compared to continuous breeding is merely the period over which conditions are suitable for offspring development and survival, there is no adaptive reason why egg or clutch sizes would be different. However, longer breeding periods may allow for an increase in the number of clutches produced during a single breeding season or year, such as that recorded in *L. fletcheri* (J. Gould, unpublished data, 2019) and *Crinia signifera* (Bull & Shine 1979; Lemckert & Shine 1993), though there is no evidence to suggest that this capacity to multi-clutch influences average clutch size among amphibians. Continuous breeding is also suggestive of species being exposed to stable conditions throughout the year where there would possibly be an opportunity to select for direct development, which would then lead to a larger egg size.

## Conclusion

Among the Australian anurans, we demonstrate for the first time that there is a clear inverse relationship between egg and clutch size, as well as positive relationships between both these variables and female body size. Such results are predicted based on the known resource limitations and physical constraints imposed upon the female during reproduction. Further, we have also shown that the evolution of an optimal clutch type is related to numerous variables pertaining to a specie’s life history and environment, in a predictable manner that is in accordance with current hypotheses regarding known reproductive trade-offs. In particular, variables are related to egg and clutch size in opposing manners, with small clutches of larger eggs more commonly observed in species that oviposit terrestrially, show direct development and possess high levels of parental care; all features of species with life histories that are not of the generalised, ancestral mode. While the variables analysed in this study are not exhaustive, they show the potential benefit of conducting big picture analyses in order to better understand the complex interdependencies that exist between a specie’s reproductive traits and their environment. Our correlation-based findings should therefore be used as a springboard to complete additional analyses on the evolution of egg and clutch sizes in this group as more detailed natural history data continues to be described in the literature. Given the comprehensiveness of this dataset and the unique biogeographical regions it covers, we therefore recommend that it is utilised for future investigations on the drivers of amphibian evolution.

